# Rapid activation of dormant type IV pili enables a dispersal–infection tradeoff in environments with fluctuating nutrients

**DOI:** 10.64898/2026.04.23.720421

**Authors:** Ahmed O. Yusuf, Zil K. Modi, Matthias D. Koch

## Abstract

Bacteria in fluctuating environments must balance the high metabolic costs of motility against risks of bacteriophage predation and immune clearance. While flagellar trade-off mechanisms are well-documented, regulation of type IV pilus (T4P) activity during environmental transitions remains unclear. We show that *Pseudomonas aeruginosa* uses an energy-dependent idling strategy to synchronize T4P-mediated surface motility with nutrient availability. In nutrient-depleted stationary phase, T4P transcription and protein levels remain constant, pre-assembled machines persist at the cell pole, yet cells produce only sparse, truncated pili that extend and retract slowly. Using a single-cell ATP biosensor, we show that T4P dynamics respond directly to cellular adenylate energy charge. Carbon source addition rapidly elevates intracellular ATP, reactivating pre-assembled T4P within minutes. This bypasses de novo protein synthesis, restoring pilus number, length, and extension/retraction rates. This rapid response drives opportunistic biofilm dispersal but, at the same time, creates an immediate tradeoff: reactivated T4P restore susceptibility to pilus-specific phages upon nutrient upshift. Thus, energetic gating of T4P enables *P. aeruginosa* to minimize exposure to phages during starvation while remaining poised for rapid reactivation. Importantly, T4P promote resistance to opsonization and phagocytosis by macrophages and neutrophils. Upon nutrient upshift, full T4P activity therefore supports dispersal and host colonization while conferring immune protection, revealing a fundamental dispersal–infection tradeoff at the host– microbe interface in fluctuating environments such as the lung and gut.

## Introduction

Type IV pili (T4P) are dynamic, retractable appendages that serve as key virulence factors in numerous clinically important pathogens, including *Pseudomonas, Neisseria*, and *Vibrio* species [1–5]. These extracellular filaments mediate critical functions such as surface attachment, twitching motility, biofilm formation, and DNA uptake through repeated cycles of extension and retraction powered by dedicated ATPases [6–10]. Upon surface attachment, retraction of a surface bound pilus generates strong mechanical force that pulls the cell forward in a process known as twitching motility [6–10]. The same mechanical interactions also enable surface sensing that regulates cyclic AMP and cyclic di-GMP (c-di-GMP) signaling pathways, thereby coupling physical contact to downstream virulence programs [11–14].

T4P play essential roles in biofilm development in vitro and during infection [15–17]. Although mutants lacking pilus assembly or retraction show clear defects in biofilm formation, the precise molecular mechanisms by which dynamic T4P cycles shape biofilm architecture remain incompletely understood. Paradoxically, functional T4P also render cells vulnerable: many bacteriophages exploit T4P as primary receptors, using pilus retraction as an entry route into the cell [18–22]. In response, some phages encode proteins that disrupt T4P function to block superinfection [23–26].

Bacteria frequently deploy multiple specialized T4P systems tailored to distinct environmental contexts. *Vibrio cholerae*, for example, expresses at least four distinct systems: the toxin-coregulated pilus (TCP), competence pili (CP), the mannose-sensitive hemagglutinin (MSHA) pilus, and the chitin-regulated pilus system (ChiRP), each optimized for specific functions such as host colonization, natural transformation, or surface attachment [1, 5]. In striking contrast, the opportunistic pathogen *Pseudomonas aeruginosa* relies on a single, highly versatile T4P system to mediate multiple behaviors simultaneously, making it an ideal model for dissecting how one machinery balances tradeoffs between conflicting demands [27].

The *P. aeruginosa* T4P machine spans the cell envelope and consists of a core assembly complex (PilMNOPQ), an inner-membrane platform (PilC), and three motor ATPases: PilB for pilus extension and PilT/PilU for retraction [1, 27–30]. The dynamic interaction of these motors with the machine complex polymerizes and depolymerizes PilA subunits from the inner membrane into the pilus fiber and controls the number and length of surface pili [4, 31–36].

Bacterial habitats are inherently fluctuating. Nutrient availability shifts rapidly in liquid environments due to flow and convection, in the host gut as digest transits, and on fomites during transmission between surfaces [37–42]. While T4P gene expression is known to respond to nutrient availability [37, 43–48] and growth-phase transitions [49–52], far less is understood about how such fluctuations directly alter T4P dynamics (e.g., pilus number, length, and activity) and, in turn, influence T4P-dependent behaviors such as biofilm formation or dispersal and phage susceptibility. This gap is particularly consequential because T4P are central to the pathogenesis of many bacteria, yet most studies have focused on transcriptional or second-messenger regulation rather than direct physical constraints imposed by cellular energy status.

Here, we address how energy-limiting conditions during nutrient transitions reshape T4P dynamics in *P. aeruginosa* and thereby modulate two key virulence traits: biofilm formation/dispersal and susceptibility to pilus-dependent bacteriophages. Together, our findings reveal how direct coupling of T4P activity to adenylate energy charge allows bacteria to remain “cloaked” from phage exploitation during starvation while remaining poised for opportunistic dispersal when conditions improve, highlighting a fundamental tradeoff between motility, dispersal, and predator avoidance in fluctuating environments.

## Results

### Pilus extension is rapidly activated upon back-dilution of stationary phase cells

To better understand how T4P dynamics change when cells transition between growth phases, for example due to nutrient shifts, we thought to initially compare the activity of T4P in stationary phase (SP) cells to exponential phase (EP) cells using Cysteine-maleimide based fluorescent labeling of the major pilin PilA to visualize pili [31, 32, 53]. Consistent with prior reports, EP cells made several active pili on most of their poles [32] (Figure 1A, Supplementary Movie 1). In stark contrast, the activity of T4P in SP cells appeared much reduced with most cells making no or only few short pili compared to EP cells (Figure 1A, Supplementary Movie 2). To verify this observation, we quantified the number of cells that extended at least one active pilus fiber in a 30 second time window (Figure 1B). Interestingly, and in support of our observation, we found that only ~ 15 % of SP cells make active pili. This ratio increased to almost 100 % within 30 minutes of transitioning the cells to fresh LB medium and remained constant over a five-hour incubation period simulating the transition from SP to EP (OD_600_ ~ 1.0).

**Figure 1.**
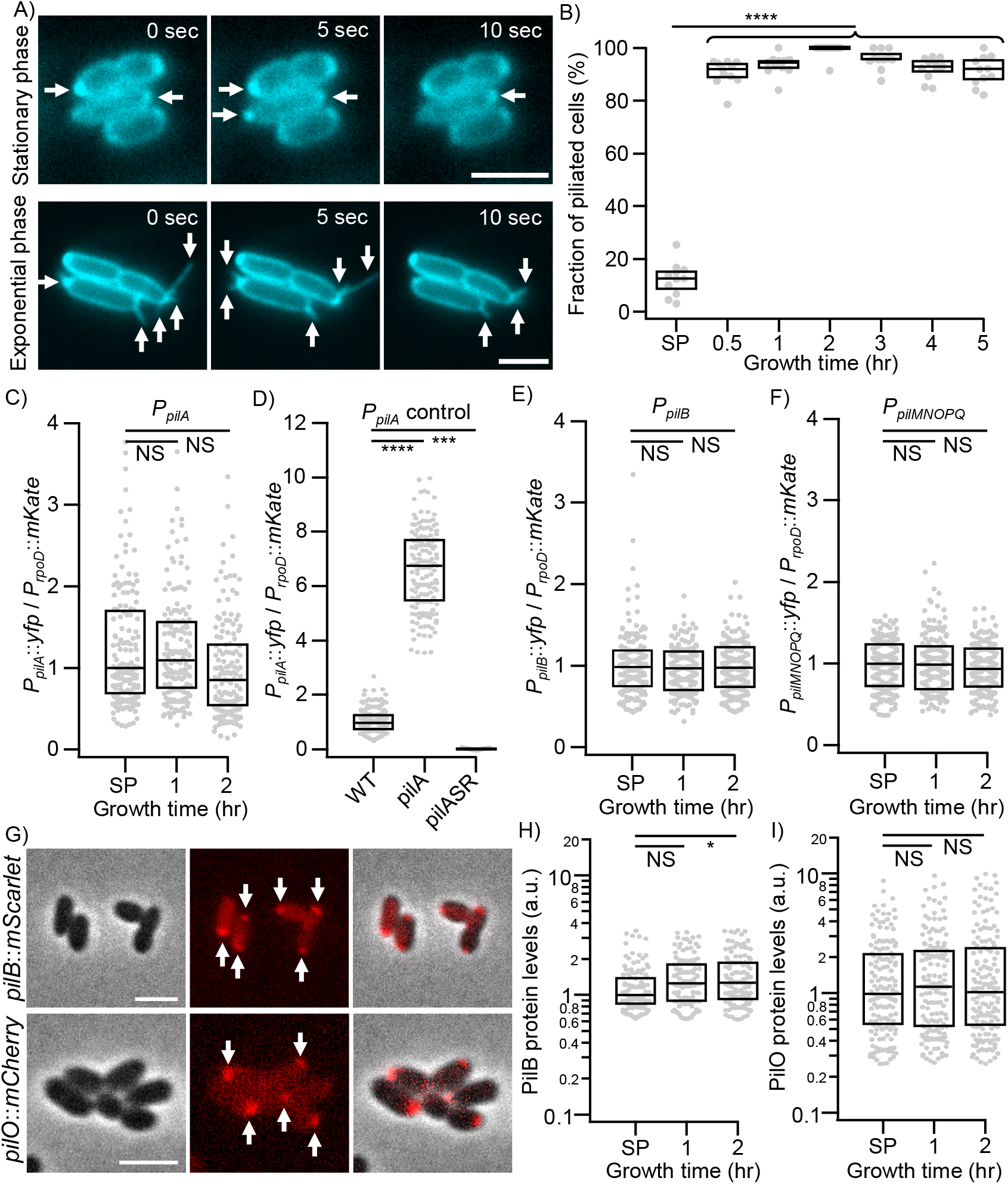
T4P machines are present in stationary phase cells and are rapidly activated in fresh medium. **A)** Time series of pilus dynamics in stationary phase (SP) and exponential phase (EP) cells. **B)** The fraction of cells that made pili in a 30 second time window of three biological and four technical replicates each. **C-F)** Transcriptional activity of the major T4P operons for the major pilin *pilA* (C,D), the extension motor *pilB* (E), and the machine scaffold operon *pilMNOPQ* (E) as a function of growth time (C,E,F) and mutant controls (D) for 50 cells of three biological replicates each. **G)** Images of protein fusions to the extension motor *pilB*::*mScarlet*-I3 and the scaffold protein *pilO*::*mCherry* in stationary phase cells. **H**,**I)** Quantification of total protein levels in individual cells for the fusions show in G for 50 cells of three biological replicates each. **Statistical analysis:** Statistical significance was tested using a bootstrapped (10,000 iterations) non-parametric Wilcoxon-Mann-Whitney two-sample rank test: (NS) not significant, P > 0.05; (*) P < 0.05; (**) P < 0.01; (***) P < 0.001; (****) P < 0.0001. **Box plots:** Boxes represent median and interquartile range (25th–75th percentiles).

### T4P machines and motors are present in stationary phase cells

The observation that cells activate pilus extension in just 30 minutes after back-dilution of SP cells into fresh growth medium made us wonder if SP cells have functional TFP machines assembled or if transcription of T4P genes is required to be rapidly activated after nutrient addition to generate machines that can extend functional pilus fibers. To test this question, we first quantified the expression of several key T4P operons using single cell fluorescent transcriptional reporters (Figure 1C-F). These reporters were made based on the established PaQa reporter that uses a constitutive *rpoD*::*mKate*2 as a control for cell growth, metabolism, and fluorophore degradation, and flipped the promoter of the *P*_*PaQa*_::*yfp* component in this reporter with the promoters for the major pilin *pilA*, the extension motor *pilB*, and the core machine complex operon *pilMNOPQ*. To our surprise, transcription of all three genes/operons remained approximately similar between SP cells, and cells after one and two hours of back-dilution. To test if our reporter constructs worked properly, we confirmed that the transcriptional activity of the *pilA* promoter is elevated in a *pilA* background and diminished in a *pilSR* background, as expected [54] (Figure 1D). Next, we tested if the T4P components are present in SP cells using fluorescent fusions PilB::mScarlet-I3 and PilO::mCherry to image cells. In support that SP cells have T4P assembly complexes present, we found bright fluorescence at the pole of SP and EP cells for both fusion proteins (Figure 1G). When we quantified changes in protein levels as a function of cell growth by measuring the total brightness of individual cell poles, we did not observe biologically significant changes between SP cells and the first two hours after back-dilution into fresh medium (Figure 1H-I). This supports our result that transcription of the T4P genes does not change significantly during transition from SP to EP and demonstrates that SP cells have T4P complexes assembled at the pole, but that these complexes are not extending pilus fibers.

### T4P become more active as cells recover from stationary phase

To understand how the T4P machines that are present in SP cells are rapidly activated to extend pilus fibers, we sought to analyze the entire dynamic extension and retraction cycles of individual pilus filaments in detail. When we compared the number of pilus fibers that individual cells made in a 30 second time window, we found that the majority of active cells made only one pilus, with few exceptions that made two or three pili (Figure 2A). Pilus activity rapidly increased to an average of two to three pili per cell 30 minutes after back-dilution and to a broad distribution of two to five pili after two hours (Figure 2B-C). Similarly, we saw the median length of pili increase from ~200 nm (SP cells) to ~900 nm (two hours back-diluted) and the maximum length increase about 4-fold from less than 1 µm to more than 3 µm (Figure 2D). Interestingly, while the median and maximum lengths increased gradually during the first two hours post back-dilution, they also decreased gradually between two and five hours post back-dilution, but remained significantly above SP levels (see Supplementary Figure 1).

**Figure 2.**
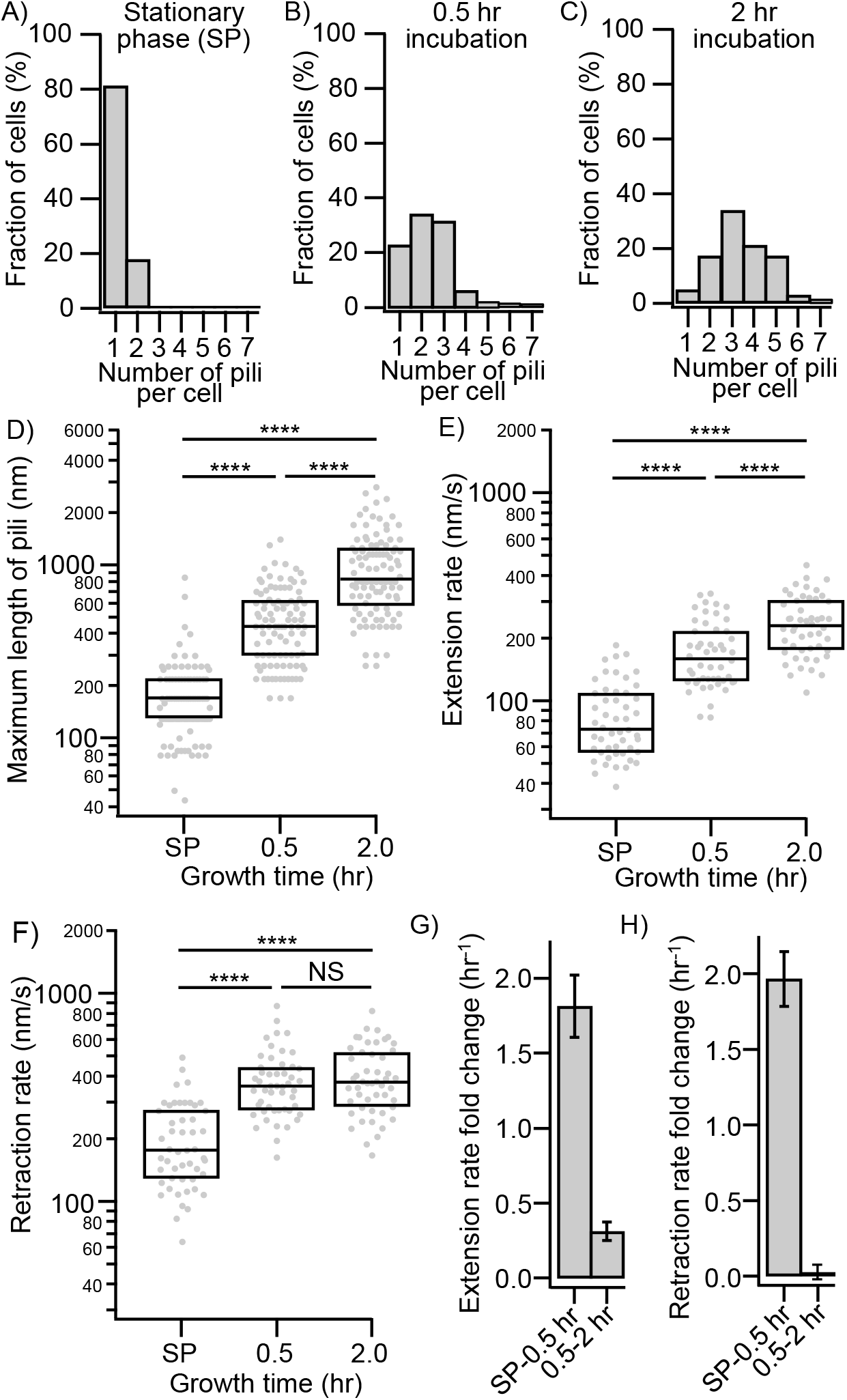
Pilus extension and retraction rates are rapidly upshifted within 30 minutes in fresh medium. **A-C)** Histograms of how many pili individual cells extended in stationary phase (SP) cells (A), and for cells incubated in fresh medium for 30 minutes (B) and 2 hours (C) for 50 cells of three biological replicates each. **D)** The maximum length of all individual pili extended by 50 cells of three biological replicates each. **E**,**F)** The extension rates (E) and retraction rates (F) of 50 individual pili of three biological replicates each. **G**,**H)** Relative fold change of the extensions rate (G) and retraction rate (H) between different growth times. **Statistical analysis:** Statistical significance was tested using a bootstrapped (10,000 iterations) non-parametric Wilcoxon-Mann-Whitney two-sample rank test: (NS) not significant, P > 0.05; (*) P < 0.05; (**) P < 0.01; (***) P < 0.001; (****) P < 0.0001. **Box plots:** Boxes represent median and interquartile range (25th–75th percentiles).

### Pili of stationary phase cells extend and retract much slower than exponential phase cells

The observation that pilus length changes dramatically as cells transition between different growth phases made us examine the rate of monomer incorporation into the fiber, given by the extension rate of individual pili (Figure 2E). SP cells extended pili slowly with a median of ~80 nm/s. This value steadily increased to ~200 nm/s at the two hour mark. This behavior is consistent with our observations for the change in pilus fiber length and suggests that pili are shorter in SP cells because they extend more slowly. Interestingly, we observed a similar trend for the retraction rate of individual pili, that increased from ~180 nm/s (SP) to 400 nm/s (2 hrs) (Figure 2F). Quantification of the relative change in extension and retraction rates between SP and 0.5 hours and 0.5 hours and 2 hours of growth shows that these rates both change about two-fold in the first 30 minutes and relatively little afterwards (Figure 2G-H). Since these processes are facilitated by ATPases (PilB and PilT), this suggests that SP cells might extend pili more slowly due to nutrient limitations, specifically due to limitation of the cellular ATP pool.

### Stationary phase cells are ATP limited and energy supplementation rapidly activates dormant T4P machines

We sought to test this hypothesis that SP cells make fewer and shorter pili because they are ATP limited by supplementing SP cells with different carbon sources directly added to spent LB (overnight culture). In support of the ATP limitation hypothesis, supplementation with either glucose, lactate, or succinate all increased the ratio of cells that made pili within 20 minutes to ~60% (Figure 3A). Consistent with the idea that the availability of these carbon sources increased cellular ATP levels, individual deletions of the three importers *oprB, dctPQM*, or *lldD* did not show a similar increase in the ration of T4P active cells. Similarly, the median and maximum lengths of pili increased from SP levels (180 nm) to levels similar of cells 30 min post back-dilution into fresh LB (~400 nm) in the same time frame that was dependent on the presence of the specific carbon source importer (Figure 3B). Together this indicates that the availability and import of nutrients rapidly triggers the activation of pilus extension.

**Figure 3.**
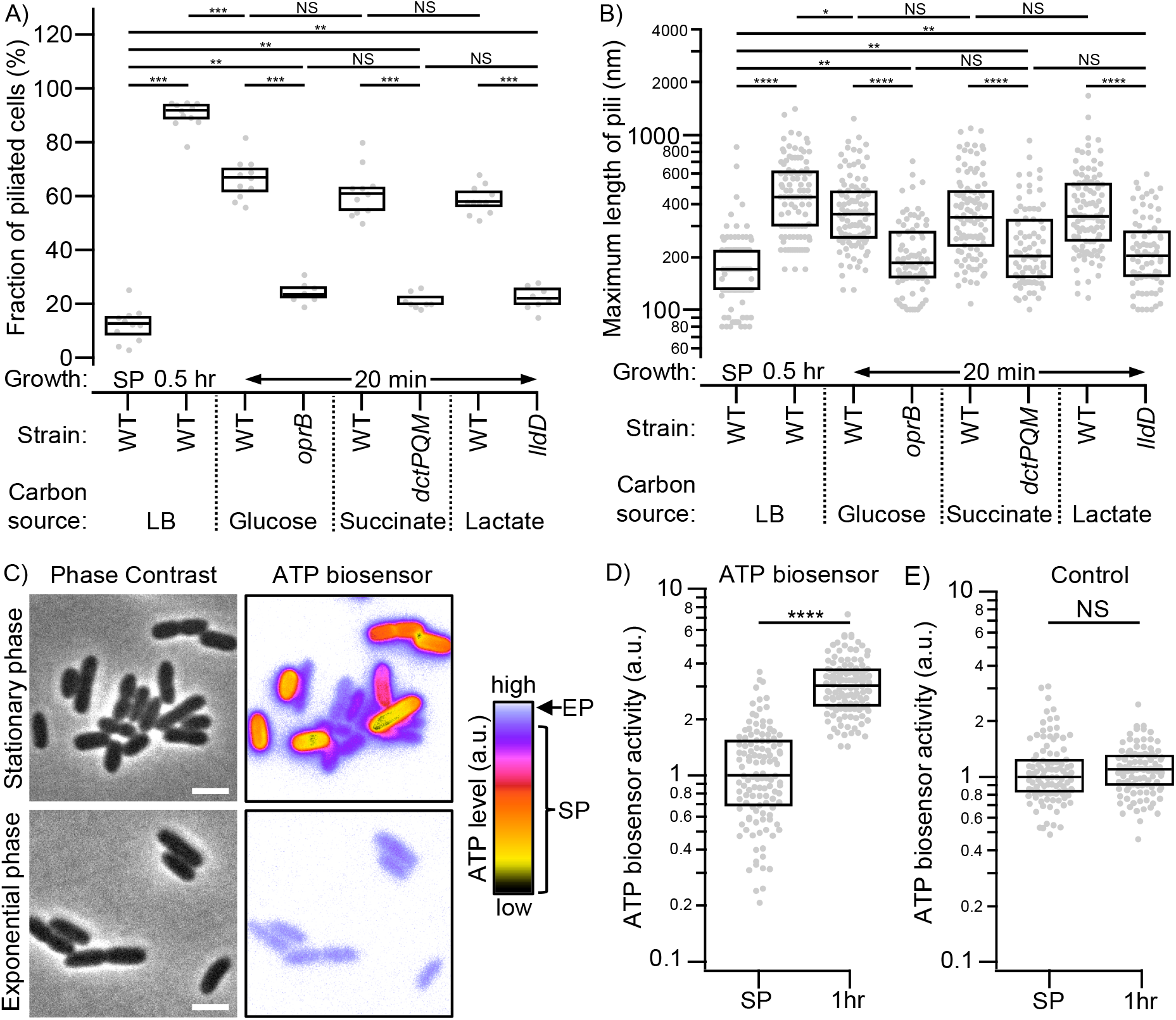
Carbon source supplementation activates T4P by increasing ATP production. **A,B)** Supplementation of stationary phase (SP) cells with different carbon sources increase the fraction of pilus producing cells (A) and pilus length (B) within 20 minutes, similar to transitioning cells to fresh rich growth medium (LB), for three biological replicates and 4 technical replicates each (A) or 50 pili for three biological replicates each (B). **C)** Images of ATP biosensor activity in individual SP and exponential phase (EP) cells. **D**,**E)** ATP biosensor activity in individual SP cells and cells grown for one hour in fresh LB medium (D) and control of a catalytically dead sensor version (E). **Statistical analysis:** Statistical significance was tested using a bootstrapped (10,000 iterations) non-parametric Wilcoxon-Mann-Whitney two-sample rank test: (NS) not significant, P > 0.05; (*) P < 0.05; (**) P < 0.01; (***) P < 0.001; (****) P < 0.0001. **Box plots:** Boxes represent median and interquartile range (25th–75th percentiles).

To determine if the supplementation of nutrients indeed yields changes in cellular ATP levels that could explain our pilus dynamics observation, we employed a fluorescent ATP reporter to measure ATP levels in single cells [55] (Figure 3C). In EP cells, the brightness of the ATP biosensor was homogenous between individual cells and cells mostly appeared dim, indicative of high ATP levels. In contrast, in SP cells, we observed a large heterogeneity between cells, with most cells expressing a bright biosensor (indicative of low ATP levels) but some cells remained a dim sensor. This is indicative of different ATP levels in a population of SP cells. We specifically quantified the ATP biosensor brightness and compared their distributions between SP cells and cells grown for one and two hours post back-dilution. (Figure 3D) These data confirm our initial assessment that SP cells are energy depleted and cellular ATP levels are increased ~3-fold when transitioning to EP. Importantly, a catalytically dead biosensor control did not change its brightness (Figure 3E). Together, this suggests that SP cells make slowly extending pili because they are ATP limited, and that supplementation of fresh nutrients activate pili because ATP levels are refueled rapidly.

### Rapid activation of dormant T4P activates biofilm dissemination and increases phage susceptibility

Next, we sought to test if rapid activation of pilus dynamics in SP cells has a functional consequence for the behavior of cells. Our findings indicate that T4P are dormant in nutrient depleted environments and activated quickly when nutrients become available again. This situation resembles liquid environments where fluid flow changes nutrient availability continuously. Here, cells accumulate into biofilms when nutrients are poor and disseminate to opportunistically explore the surroundings when conditions are favorable. We wondered if the activation of T4P supports this behavior and analyzed the dispersal of cells from biofilms after we added glucose to biofilms in SP. We first grew biofilms in a standard biofilm assay on pegs submerged into growth medium in 96 well plate format using WT cells and a pilus null mutant (*pilA*) that is unable to assemble pilus filaments [56]. After biofilms had formed and cells reached SP, we treated the peg-associated biofilms either with glucose or a vehicle control (MOPS buffer), stained the residual biofilm with crystal violet, and subsequently dissolved and quantified the dye in a plate reader by OD_600_ measurement. Consistent with previous research, we observed that both WT and *pilA* were able to form robust biofilms (untreated) (Figure 4A). After treatment of WT cells, the amount of residual biofilm was significantly larger (~50%) in the vehicle control compared to the glucose treated environment, and the amount of biofilm dropped significantly compared to the untreated condition in both cases (Figure 4B). In support of our hypothesis that this difference is due to the activation of T4P, we found no significant difference between both treatments in the *pilA* mutant. Comparing WT cells to *pilA*, we did see a large decrease in residual biofilm with and without treatment, indicating that even low levels of pilus activity (as in SP cells) significantly support dispersal of cells from mature biofilms and that more active T4P increase dispersal.

**Figure 4.**
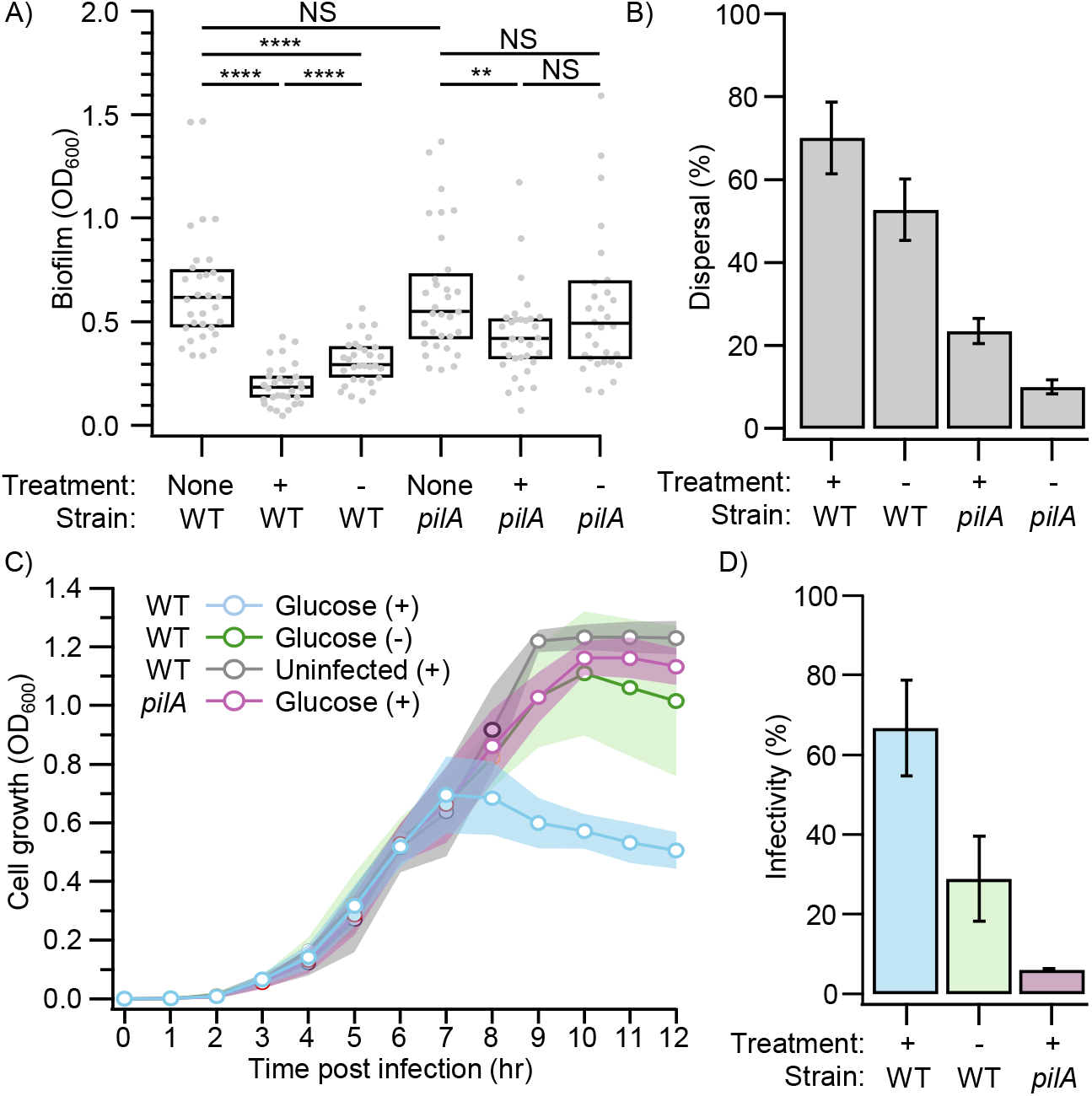
Activation of dormant T4P by nutrient upshift triggers biofilm dispersal and increases susceptibility to phage infection. **A)** Biofilm biomass before treatment (treatment: none) and after treatment with (+) and without (-) glucose in WT cells capable of extending pili and a *pilA* mutant control unable to extend pili. **B)** Dispersal measured by the decrease in biofilm biomass relative to the no treatment control. **C)** Growth of WT cells (un-)infected with T4P specific phage JBD68 with (+) and without (-) glucose and a *pilA* control. **D)** Infectivity measured by the decrease in cell growth at the last time point relative to the uninfected control. **Statistical analysis:** Statistical significance was tested using a bootstrapped (10,000 iterations) non-parametric Wilcoxon-Mann-Whitney two-sample rank test: (NS) not significant, P > 0.05; (*) P < 0.05; (**) P < 0.01; (***) P < 0.001; (****) P < 0.0001. **Box plots:** Boxes represent median and interquartile range (25th–75th percentiles). **Error bars:** represent the mean and standard deviation.

This result that the activation of pili supports the dissemination of biofilms posed the question if this creates a tradeoff with infection. As T4P become more dynamic and extend longer filaments, they might also become more susceptible to bacteriophages. We tested this idea by monitoring cell growth of SP cells after 30 minutes of glucose incubation (to activate pili) that were then subjected to the pilus specific phage JBD68 for two minutes (to infect cells). Both uninfected cells and the *pilA* mutant grew to SP at high OD ~ 1.2, as expected (Figure 4C). Infected WT cells displayed reduced growth due to cell lysis in a glucose-dependent manner. The no glucose control showed only a small reduction in growth (due to low activity of T4P) while the glucose treated cells had a substantial reduction in growth by ~ 70% compared to the uninfected control (Figure 4D). This demonstrates that the increase in activity of T4P exposes cells to the risk of phage infection (Figure 5).

**Figure 5.**
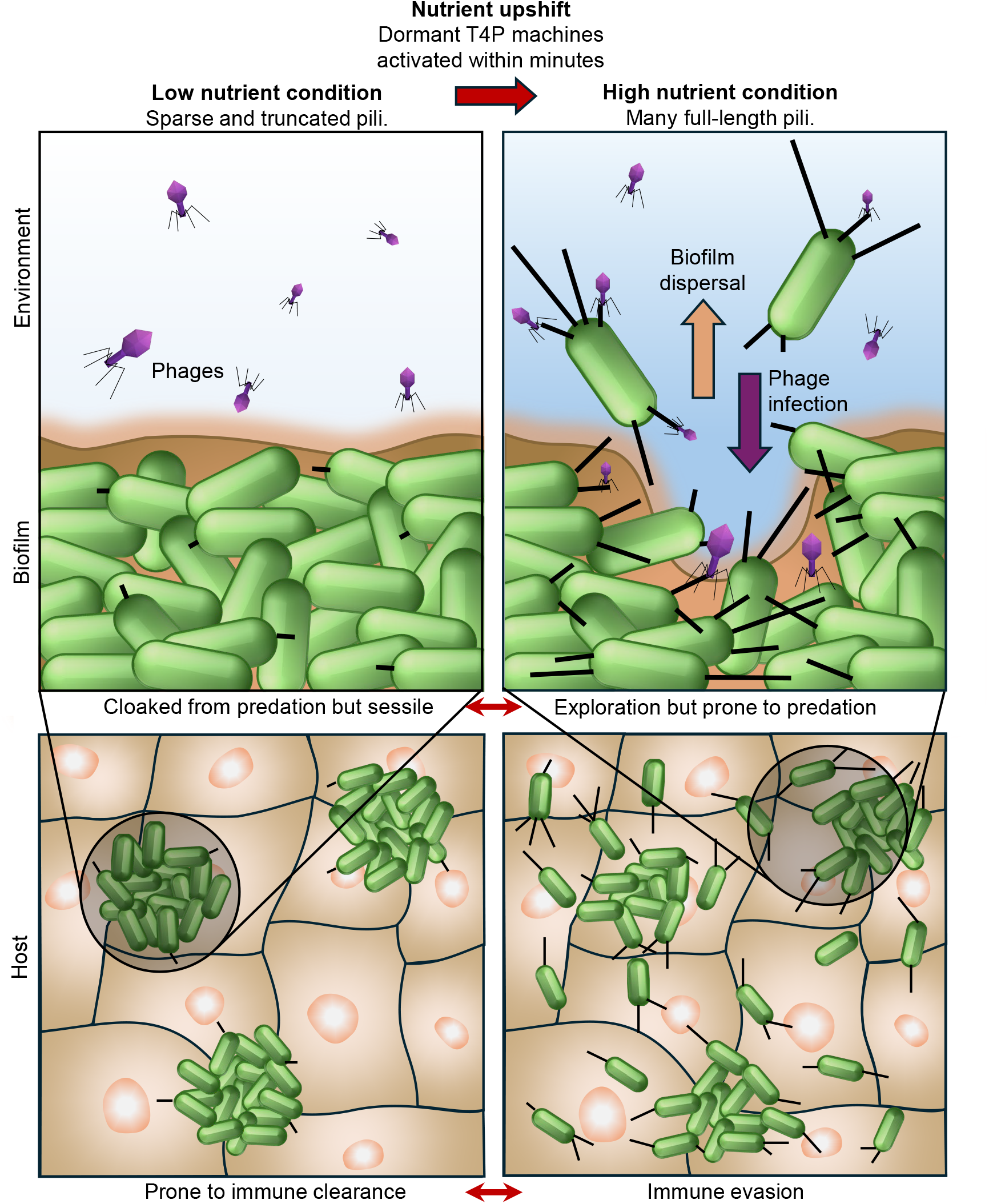
Schematic of how reactivation of dormant T4P upon nutrient upshift presents a tradeoff between dispersal and predation by the host or bacteriophages in the environment.

## Discussion

Bacteria use motility to opportunistically explore environments, for example to find nutrient patches or to colonize new niches in fluctuating host environments like the gut or lung. This behavior comes at the expense of substantial metabolic costs and increased risks of immune clearance or phage infection. How bacteria strike this balance has been well worked out for flagella-based motility. Construction and operation of flagella consume roughly 10% of a cell’s total energy budget in *E. coli* and up to ~40% in some taxa, [57]. In nutrient-limited or starvation conditions, bacteria dynamically adjust motility investment in proportion to anticipated chemotactic benefit, while marine copiotrophs display a clear risk–reward dichotomy: some strains sustain costly swimming for days by converting biomass into energy to forage for new patches (limokinetic), whereas others rapidly arrest motility to conserve resources (limostatic) [58, 59]. Active flagella also heighten susceptibility to phages, as mutations conferring phage resistance frequently reduce motility through regulatory pathways such as the Rcs phosphorelay [60].

Here, we show that *Pseudomonas aeruginosa* employs an energy-dependent “idling” strategy for type IV pili (T4P)-mediated surface motility that parallels these flagellar trade-offs while using a distinct mechanistic solution. In nutrient-depleted stationary phase, T4P transcription and protein levels (PilA, PilB, PilO) remain largely constant and pre-assembled machines persist at the cell pole, yet cells produce only sparse, truncated pili that extend and retract slowly (~80 nm/s extension rate). Supplementation of carbon sources (glucose, lactate, or succinate) directly to spent medium rapidly elevates intracellular ATP levels, as monitored by a fluorescent biosensor, and activates pilus dynamics within minutes. This restores the fraction of active cells, increases pilus number and length, and accelerates extension (~80 to ~200 nm/s) and retraction rates without the need for de novo protein synthesis. Consequently, nutrient upshift triggers quick dispersal of cells from mature biofilms, evidenced by significantly lower residual biofilm biomass in WT cultures compared with *pilA* mutants after glucose addition, including greater baseline dispersal even in vehicle controls. However, this rapid activation also heightens susceptibility to pilus-specific phages like JBD68 in a glucose-dependent manner (~70% growth reduction in activated WT cells), highlighting a tradeoff between motility and infection (Figure 5).

This energy-gated idling of T4P may be particularly relevant in host environments such as the lung, where nutrient availability fluctuates locally and intact pili help *P. aeruginosa* resist opsonization and clearance by alveolar macrophages and neutrophils [61, 62]. By maintaining low T4P activity under nutrient stress, cells could minimize unnecessary surface exposure while poised for rapid reactivation that supports dispersal and colonization when conditions improve (Figure 5).

This T4P idling strategy shares important functional parallels with flagellar motility. Both systems couple dispersal benefit (biofilm escape or nutrient-finding) with an immediate infection cost, while minimizing unnecessary energy expenditure during starvation. The primary mechanistic difference lies in the mode of control: flagellar systems frequently rely on transcriptional regulation, growth-rate sensing, or strain-specific behavioral choices, whereas T4P activity here is closely linked to cellular energy status in a manner consistent with direct energetic gating of the extension (PilB) and retraction (PilT) motors. This low-latency, post-translational response may be particularly advantageous for surface-associated lifestyles in competitive or flow-variable environments [38, 63, 64].

Our biofilm dispersal data further suggest that even low-level T4P activity in stationary phase contributes to reduced biofilm robustness in WT versus *pilA* cells. While extensive work has addressed T4P roles in early surface colonization and macroscopic biofilm patterning (e.g., via cell arrangement and matrix interactions), how dynamic T4P modulation shapes local architecture or mechanical properties within established biofilms remains less explored [15, 65–70]. Recent studies show that *P. aeruginosa* biofilms exhibit highly ordered cellular arrangements, including vertical striations of lengthwise-packed cells across much of the biofilm depth, and that mutations altering surface structures (including those affecting pili) can disrupt this microstructure, influencing local metabolic activity and permeability. Whether dynamic, energy-gated changes in T4P activity fine-tune microscopic cohesion, cell packing, or mechanical integrity in mature biofilms warrants further investigation.

Overall, these findings highlight a conserved ecological logic across motility systems: costly appendages are kept in a poised but throttled state during nutrient stress to minimize metabolic burden and infection risk, then rapidly deployed when conditions improve. By linking T4P dynamics directly to adenylate energy charge, *P. aeruginosa* achieves near-instantaneous biofilm escape without heavy biosynthetic costs. This strategy could enable opportunistic pathogens to navigate the complex risk–reward decisions required for persistence and dissemination at the host–microbe interface [71, 72]. Future experiments will test whether similar energetic idling occurs in other T4P- or flagella-bearing species. This strategy may represent a widespread adaptation that allows bacteria to protect the population under starvation while exploiting transient opportunities for dispersal and colonization.

## Acknowledgements

We would like to thank Sayak Muckhopadhyay and Pushkar Lele for the recommendation of the ATP biosensor and stimulating discussion and Josh Brehm and Joe Sorg for suggesting and lending the Cerillo plate reader. We would like to thank the entire Koch lab for stimulating discussion and the Department of Biology at Texas A&M for its supportive environment.

This work was supported by grant R35GM155280 from the National Institute of Health and startup funds from the College of Arts and Sciences, Division of Research, and Department of Biology at Texas A&M University to M.D.K.

## Author contributions

A.O.Y and M.D.K. designed research and wrote the manuscript. A.O.Y and Z.K.M provided reagents, genetic constructs, and performed experiments. A.O.Y, Z.K.M., and M.D.K. analyzed data.

## Methods and Protocols

### Strains and Growth Conditions

Information on cloning, plasmids, and primers used in this study as well as detailed methods on individual assays can be found in the Supplementary Information (SI) Methods and Protocols.

*P. aeruginosa* PAO1 was grown in liquid lysogeny broth (LB) Miller (Difco) and EZ rich defined medium (Teknova) in a floor shaker, on LB Miller agar (1.5 % Bacto Agar), on Vogel-Bonner minimal medium (VBMM) agar (200 mg/l MgSO4 7H2O, 2 g/l citric acid, 10 g/l K2HPO4, 3.5 g/l NaNH4HPO4 4 H2O, and 1.5% agar), and on no-salt LB (NSLB) agar (10 g/l tryptone, 5 g/l yeast extract, and 1.5% agar) at 30 °C (for cloning) or at 37 °C. *Escherichia coli* S17 was grown in liquid LB Miller (Difco) in a floor shaker and on LB Miller agar (1.5 % Bacto Agar) at 30 °C (for cloning, SI Appendix) or at 37 °C. Antibiotics were used at the following concentrations: 200 μg/mL carbenicillin in liquid (300 μg/mL on plates) or 10 μg/mL gentamycin in liquid (30 μg/mL on plates) in liquid for *P. aeruginosa*, and 100 μg/mL carbenicillin in liquid (100 μg/mL on plates) or 30 μg/mL gentamycin in liquid (30 μg/mL on plates) for *E. coli*.

## References

1. Ellison, C.K., G.B. Whitfield, and Y.V. Brun, Type IV Pili: dynamic bacterial nanomachines. FEMS Microbiol Rev, 2022. 46(2).

2. Hoppe, J., et al., PilY1 Promotes Legionella pneumophila Infection of Human Lung Tissue Explants and Contributes to Bacterial Adhesion, Host Cell Invasion, and Twitching Motility. Front Cell Infect Microbiol, 2017. 7: p. 63.

3. Floyd, K.A., et al., c-di-GMP modulates type IV MSHA pilus retraction and surface attachment in Vibrio cholerae. Nature Communications, 2020. 11(1): p. 1549.

4. Ellison, C.K., et al., A bifunctional ATPase drives tad pilus extension and retraction. Science Advances, 2019. 5(12): p. eaay2591.

5. Ellison, C.K., et al., Subcellular localization of type IV pili regulates bacterial multicellular development. Nature Communications, 2022. 13(1): p. 6334.

6. Henrichsen, J., Twitching motility. Annu Rev Microbiol, 1983. 37: p. 81–93.

7. Siryaporn, A., et al., Colonization, competition, and dispersal of pathogens in fluid flow networks. Current Biology, 2015. 25(9): p. 1201–1207.

8. Shen, Y., et al., Flow directs surface-attached bacteria to twitch upstream. Biophys J, 2012. 103(1): p. 146–51.

9. Oliveira, N.M., K.R. Foster, and W.M. Durham, Single-cell twitching chemotaxis in developing biofilms. Proceedings of the National Academy of Sciences, 2016. 113(23): p. 6532–6537.

10. Jin, F., et al., Bacteria use type-IV pili to slingshot on surfaces. Proceedings of the National Academy of Sciences, 2011. 108(31): p. 12617–12622.

11. Koch, M.D., et al., <i>Pseudomonas aeruginosa</i> distinguishes surfaces by stiffness using retraction of type IV pili. Proceedings of the National Academy of Sciences, 2022. 119(20): p. e2119434119.

12. Luo, Y., et al., A Hierarchical Cascade of Second Messengers Regulates Pseudomonas aeruginosa Surface Behaviors. mBio, 2015. 6(1): p. 10.1128/mbio.02456-14.

13. Persat, A., et al., Type IV pili mechanochemically regulate virulence factors in Pseudomonas aeruginosa. Proc Natl Acad Sci U S A, 2015. 112(24): p. 7563–8.

14. Siryaporn, A., et al., Surface attachment induces <i>Pseudomonas aeruginosa</i> virulence. Proceedings of the National Academy of Sciences, 2014. 111(47): p. 16860–16865.

15. Klausen, M., et al., Biofilm formation by Pseudomonas aeruginosa wild type, flagella and type IV pili mutants. Mol Microbiol, 2003. 48(6): p. 1511–24.

16. Haagensen, J.A.J., et al., Differentiation and Distribution of Colistin- and Sodium Dodecyl Sulfate-Tolerant Cells in <i>Pseudomonas aeruginosa</i> Biofilms. Journal of Bacteriology, 2007. 189(1): p. 28–37.

17. Chiang, P. and L.L. Burrows, Biofilm formation by hyperpiliated mutants of Pseudomonas aeruginosa. J Bacteriol, 2003. 185(7): p. 2374–8.

18. Meng, R., et al., Structural basis of Acinetobacter type IV pili targeting by an RNA virus. Nature Communications, 2024. 15(1): p. 2746.

19. Wang, Y., et al., Structural mechanisms of Tad pilus assembly and its interaction with an RNA virus. Science Advances, 2024. 10(18): p. eadl4450.

20. Harvey, H., et al., Pseudomonas aeruginosa defends against phages through type IV pilus glycosylation. Nature Microbiology, 2018. 3(1): p. 47–52.

21. Bae, H.-W., et al., Pilin regions that select for the small RNA phages in <i>Pseudomonas aeruginosa</i> type IV pilus. Journal of Virology, 2025. 99(4): p. e01949–24.

22. Chung, I.-Y., et al., A phage protein that inhibits the bacterial ATPase required for type IV pilus assembly. Proceedings of the National Academy of Sciences, 2014. 111(31): p. 11503–11508.

23. Hendrix, H., et al., PlzR regulates type IV pili assembly in Pseudomonas aeruginosa via PilZ binding. Nature Communications, 2024. 15(1): p. 8717.

24. Taylor, V.L., et al., Prophages block cell surface receptors to preserve their viral progeny. Nature, 2025. 644(8078): p. 1049–1057.

25. Schmidt, A.K., et al., A Filamentous Bacteriophage Protein Inhibits Type IV Pili To Prevent Superinfection of Pseudomonas aeruginosa. mBio, 2022. 13(1): p. e02441–21.

26. Thongchol, J., et al., Removal of <i>Pseudomonas</i> type IV pili by a small RNA virus. Science, 2024. 384(6691): p. eadl0635.

27. Burrows, L.L., Pseudomonas aeruginosa twitching motility: type IV pili in action. Annu Rev Microbiol, 2012. 66: p. 493–520.

28. Roberge, N.A. and L.L. Burrows, Building permits—control of type IV pilus assembly by PilB and its cofactors. Journal of Bacteriology, 2024. 206(12): p. e00359–24.

29. Guo, S., et al., PilY1 regulates the dynamic architecture of the type IV pilus machine in Pseudomonas aeruginosa. Nature Communications, 2024. 15(1): p. 9382.

30. Chang, Y.-W., et al., Architecture of the Type IVA Pilus Machine. Biophysical Journal, 2016. 110(3): p. 468a–469a.

31. Ellison, C.K., et al., Real-time microscopy and physical perturbation of bacterial pili using maleimide-conjugated molecules. Nature Protocols, 2019. 14(6): p. 1803–1819.

32. Koch, M.D., et al., Competitive binding of independent extension and retraction motors explains the quantitative dynamics of type IV pili. Proceedings of the National Academy of Sciences, 2021. 118(8): p. e2014926118.

33. Talà, L., et al., Pseudomonas aeruginosa orchestrates twitching motility by sequential control of type IV pili movements. Nature Microbiology, 2019. 4(5): p. 774–780.

34. Kühn, M.J., et al., Two antagonistic response regulators control Pseudomonas aeruginosa polarization during mechanotaxis. Embo j, 2023. 42(7): p. e112165.

35. Chlebek, J.L., et al., PilT and PilU are homohexameric ATPases that coordinate to retract type IVa pili. PLOS Genetics, 2019. 15(10): p. e1008448.

36. Teipen, A.E., et al., Structural modeling reveals the mechanism of motor ATPase coordination during type IV pilus retraction. bioRxiv, 2025: p. 2025.10.30.685630.

37. Padron, G.C., et al., Shear flow promotes bacterial growth and shapes spatial gradients by rapidly replenishing scarce nutrients. mBio. 0(0): p. e03446–25.

38. Shuppara, A.M., et al., Shear flow patterns antimicrobial gradients across bacterial populations. Science Advances, 2025. 11(11): p. eads5005.

39. Gilbert, J.A., et al., Current understanding of the human microbiome. Nature Medicine, 2018. 24(4): p. 392–400.

40. Kuziel, G.A. and S. Rakoff-Nahoum, The gut microbiome. Curr Biol, 2022. 32(6): p. R257–r264.

41. Stephens, B., et al., Microbial Exchange via Fomites and Implications for Human Health. Curr Pollut Rep, 2019. 5(4): p. 198–213.

42. Lopez, G.U., et al., Transfer efficiency of bacteria and viruses from porous and nonporous fomites to fingers under different relative humidity conditions. Appl Environ Microbiol, 2013. 79(18): p. 5728–34.

43. Krebs, S.J. and R.K. Taylor, Nutrient-dependent, rapid transition of Vibrio cholerae to coccoid morphology and expression of the toxin co-regulated pilus in this form. Microbiology, 2011. 157(10): p. 2942–2953.

44. Ni, L., et al., Bacteria differently deploy type-IV pili on surfaces to adapt to nutrient availability. npj Biofilms and Microbiomes, 2016. 2(1): p. 15029.

45. Edén, C.S. and H.A. Hansson, Escherichia coli pili as possible mediators of attachment to human urinary tract epithelial cells. Infection and Immunity, 1978. 21(1): p. 229–237.

46. Mokrzan, E.M., T.J. Johnson, and L.O. Bakaletz, Expression of the Nontypeable Haemophilus influenzae Type IV Pilus Is Stimulated by Coculture with Host Respiratory Tract Epithelial Cells. Infection and Immunity, 2019. 87(12): p. 10.1128/iai.00704-19.

47. Chirakadavil, J.B., et al., Calcium signals natural transformation in <em>Acinetobacter baumannii</em>. bioRxiv, 2026: p. 2026.02.23.707608.

48. Denise, R., S.S. Abby, and E.P.C. Rocha, Diversification of the type IV filament superfamily into machines for adhesion, protein secretion, DNA uptake, and motility. PLoS Biol, 2019. 17(7): p. e3000390.

49. Bradley, D.E., A study of pili on Pseudomonas aeruginosa. Genetical Research, 1972. 19(1): p. 39–51.

50. Vesel, N. and M. Blokesch, Pilus Production in Acinetobacter baumannii Is Growth Phase Dependent and Essential for Natural Transformation. Journal of Bacteriology, 2021. 203(8): p. 10.1128/jb.00034-21.

51. Song, X.-M., et al., The growth phase-dependent regulation of the pilus locus genes by two-component system TCS08 in Streptococcus pneumoniae. Microbial Pathogenesis, 2009. 46(1): p. 28–35.

52. Salzer, R., et al., Environmental factors affecting the expression of type IV pilus genes as well as piliation of Thermus thermophilus. FEMS Microbiology Letters, 2014. 357(1): p. 56–62.

53. Ellison, C.K., et al., Obstruction of pilus retraction stimulates bacterial surface sensing. Science, 2017. 358(6362): p. 535–538.

54. Kilmury, S.L. and L.L. Burrows, Type IV pilins regulate their own expression via direct intramembrane interactions with the sensor kinase PilS. Proceedings of the National Academy of Sciences, 2016. 113(21): p. 6017–6022.

55. Yaginuma, H., et al., Diversity in ATP concentrations in a single bacterial cell population revealed by quantitative single-cell imaging. Scientific Reports, 2014. 4(1): p. 6522.

56. Marko, V.A., et al., Pseudomonas aeruginosa type IV minor pilins and PilY1 regulate virulence by modulating FimS-AlgR activity. PLoS Pathog, 2018. 14(5): p. e1007074.

57. Schavemaker, P.E. and M. Lynch, Flagellar energy costs across the tree of life. elife, 2022. 11: p. e77266.

58. Ni, B., et al., Growth-rate dependent resource investment in bacterial motile behavior quantitatively follows potential benefit of chemotaxis. Proceedings of the National Academy of Sciences, 2020. 117(1): p. 595–601.

59. Keegstra, J.M., et al., Risk–reward trade-off during carbon starvation generates dichotomy in motility endurance among marine bacteria. Nature Microbiology, 2025. 10(6): p. 1393–1403.

60. Burmeister, A.R., H. Tewatia, and C. Skinner, A tradeoff between bacteriophage resistance and bacterial motility is mediated by the Rcs phosphorelay in Escherichia coli. Microbiology, 2024. 170(8): p. 001491.

61. Tan, R.M., et al., Type iv pilus of pseudomonasaeruginosa confers resistance to antimicrobial activities of the pulmonary surfactant protein-a. Journal of innate immunity, 2014. 6(2): p. 227–239.

62. Tan, R.M., et al., Type IV pilus glycosylation mediates resistance of Pseudomonas aeruginosa to opsonic activities of the pulmonary surfactant protein A. Infection and immunity, 2015. 83(4): p. 1339–1346.

63. Padron, G.C., et al., Bacteria in Fluid Flow. Journal of bacteriology, 2023. 205(4): p. e00400–22.

64. Padron, G.C., et al., Shear flow promotes bacterial growth and shapes spatial gradients by rapidly replenishing scarce nutrients. mBio, 2026: p. e03446–25.

65. Dayton, H., et al., Cellular arrangement impacts metabolic activity and antibiotic tolerance in Pseudomonas aeruginosa biofilms. Plos Biology, 2024. 22(2): p. e3002205.

66. Gomez, S., et al., Substrate stiffness impacts early biofilm formation by modulating Pseudomonas aeruginosa twitching motility. Elife, 2023. 12: p. e81112.

67. Nijjer, J., et al., Mechanical forces drive a reorientation cascade leading to biofilm self-patterning. Nature communications, 2021. 12(1): p. 6632.

68. Huang, X., et al., Vibrio cholerae biofilms use modular adhesins with glycan-targeting and nonspecific surface binding domains for colonization. Nature communications, 2023. 14(1): p. 2104.

69. Hartmann, R., et al., Emergence of three-dimensional order and structure in growing biofilms. Nature physics, 2019. 15(3): p. 251–256.

70. Vidakovic, L., et al., Biofilm formation on human immune cells is a multicellular predation strategy of Vibrio cholerae. Cell, 2023. 186(12): p. 2690–2704.e20.

71. Sahu, A. and R. Ruhal, Immune system dynamics in response to Pseudomonas aeruginosa biofilms. npj Biofilms and Microbiomes, 2025. 11(1): p. 104.

72. Swart, A.L., et al., Pseudomonas aeruginosa breaches respiratory epithelia through goblet cell invasion in a microtissue model. Nature Microbiology, 2024. 9(7): p. 1725–1737.

