## Supplementary Data and Methods for "Rapid activation of dormant type IV pili enables a dispersal–infection tradeoff in environments with fluctuating nutrients"

**Supplementary Results**

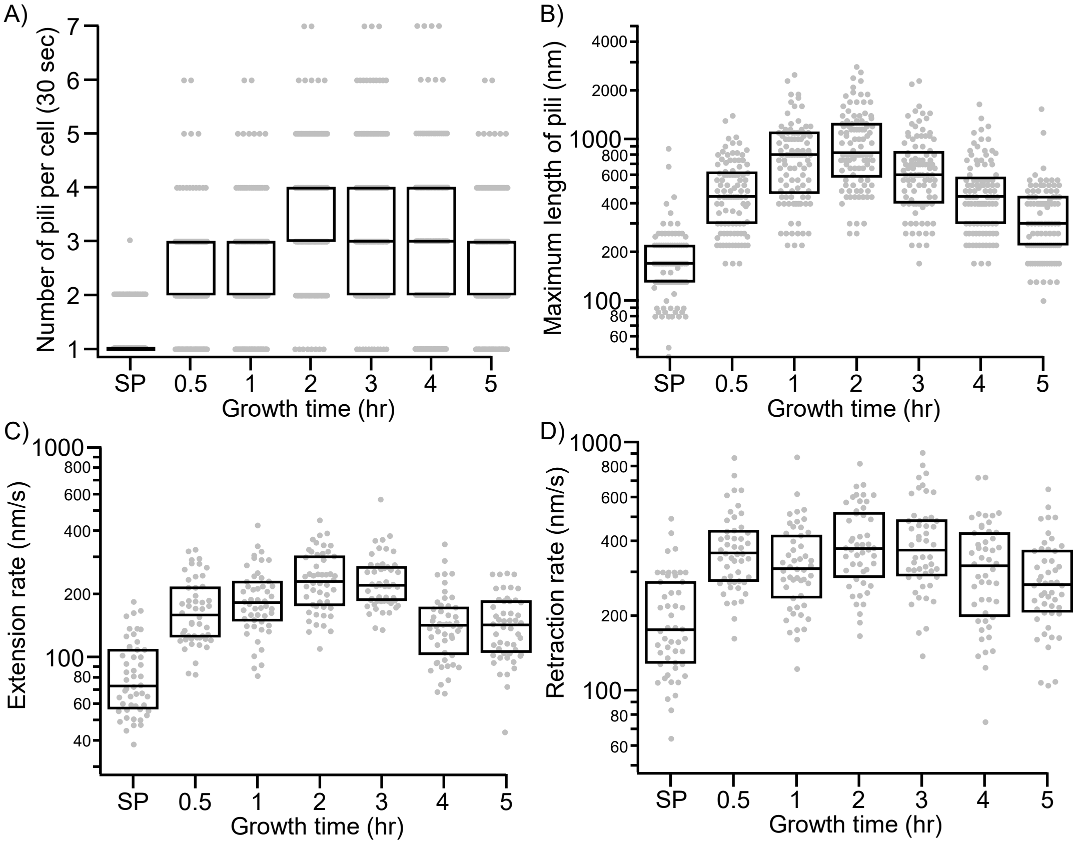


**Supplementary Figure 1. Pilus dynamics as a function of growth time. A)** The number of pili made by individual cells in a 30 second time window. Only cells that made at least one pilus were counted. **B)** The maximum length of pili. **C)** The rate of pilus extension. **D)** The rate of pilus retraction. **Statistical analysis:** Each plot shows 50 individual cells (A) or pili (B-D) of three biological replicates each**. Box plots:** Boxes represent median and interquartile range (25th–75th percentiles).

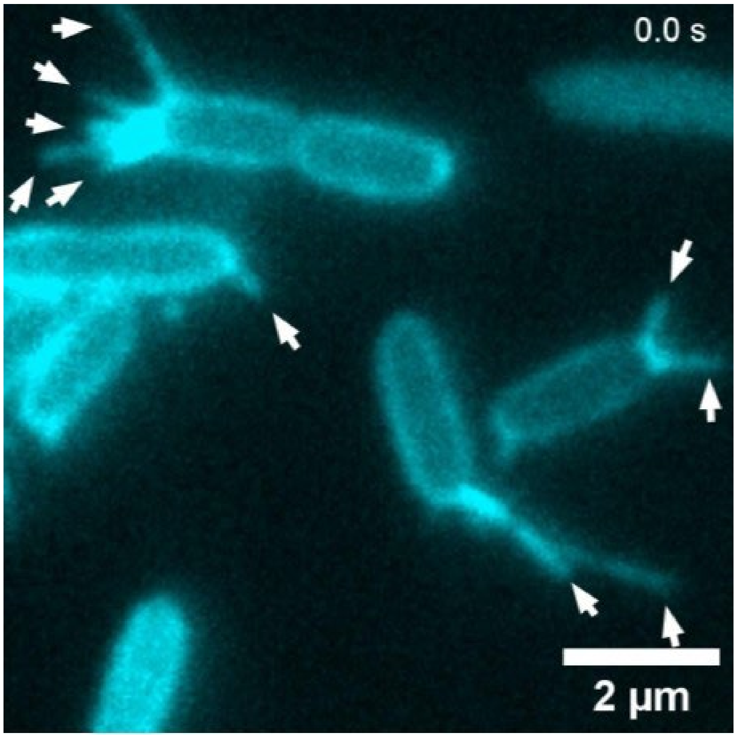


**Supplementary Movie 1. Pilus extension (white arrows) of a culture 1 hour post back dilution into fresh LB medium.**

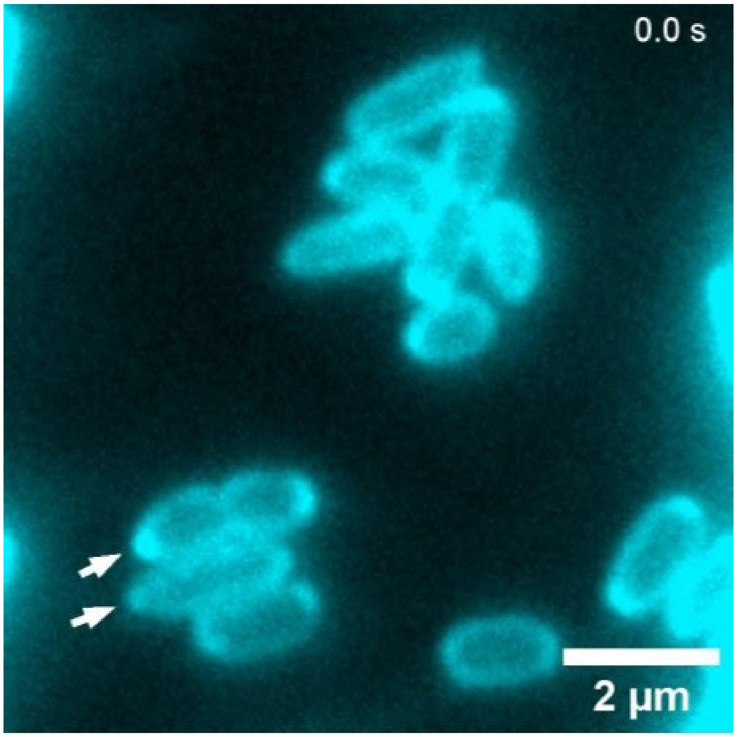


**Supplementary Movie 2. Pilus extension (white arrows) in an overnight (stationary phase, SP) cell culture.**

### Supplementary Methods and Protocols

#### Generation of chromosomal mutants (*oprB1*, *lldD*, *detPQM*)

Chromosomal mutants were generated using allelic exchange [1]. For deletion mutants (*oprB1*, *lldD*, *dctPQM*), primers for the 500 bp flanking regions upstream (P1/P2) and downstream (P3/P4) of the gene of interest were generated. The flanking regions were PCR amplified respectively and joined together in a second step using overlap extension PCR (SOE-PCR). These fragments were then cloned into the suicide vector pEXG2 (GentR) using restriction cloning. The plasmid was transformed into *E. coli* S17 cells by electroporation and plated on LB plates containing the antibiotic marker (Gent).

Similarly, a fluorescent fusion of *pilB::mScarlet*-I3 was generated by amplifying 500bp genomic fragments immediately upstream and downstream of the stop codon of *pilB* using primers pilB_FTL1_CT_F1.FOR/REV and pilB_FTL1_CT_F2.FOR/REV. The *mScarlet*-i3 fluorophore was amplified using primers FTL1_FP.FOR/REV. The fragments were joined with the pEXG2 backbone using NEBuilder HiFi DNA Assembly Master Mix (New England Biolabs) according to the manufacturer's protocol.

Successful transformation for all constructs was confirmed using colony PCR and sanger sequencing with primers pEXG2_Ver1/2. Constructs were conjugated into the recipient strain by growing 1.5 mL transformed *E. coli* cells to OD 0.5. The *Pseudomonas aeruginosa* recipient strain was grown overnight, and 0.5 ml culture was diluted 1:2 into fresh LB and incubated for 3 hours at 42 °C. Both cultures were pelleted into 100 μL and spotted onto an LB agar plate and incubated overnight at 30 °C. The puddle was scrapped off, resuspended in 150 μL PBS, then spread onto a VBMM plate containing 30 μg/ml gentamycin and incubated for 24 hours at 37 °C. Eight single colonies from the VBMM plate were struck onto NSLB and incubated for 24 hours at 30 °C. Several single colonies from the NSLB plate were screened for the correct mutation using PCR amplification with the flaking primers and sanger sequencing. All strains were stored at −80 °C in 25 % (v/v) glycerol

#### ATP biosensor plasmids construction and chromosomal integration

The pAY9 (ATP biosensor) and pAY11 (catalytic dead mutant) plasmids were constructed with modification to the established single fluorophore Queens ATP biosensor which uses a circularly permuted green fluorescent protein (cpGFP) introduced between the F0F1 ATP synthase[2]. In brief, cDNA coding for the respective amino acid sequences were ordered codon optimized for *Pseudomonas aeruginosa* from IDT. Plasmid pMK89 was linearized by digestion with AvrII and Eco53KI and all fragments were assembled using NEBuilder HiFi DNA Assembly Master Mix (New England Biolabs) according to the manufacturer’s instruction. The assembly products were electroporated into electrocompetent *E. coli* S17, then recovered in 1 mL LB and grown at 37 °C with shaking for 1.5 hour. Following recovery, cells were concentrated by centrifugation, resuspended in 100 µL LB, and plated on LB agar supplemented with gentamycin. Successful transformants were confirmed by colony PCR using primers pTN7_Ver3/4 and sanger sequencing, and stored at −80 °C in 25 % (v/v) glycerol.

The resulting plasmids were extracted from *E. coli* S17 using the Qiagen miniprep kit and transferred into recipient strains by electroporation using co-transformation of plasmid pTNS2. The recipient strain was made electrocompetent by growing the cells to late log phase and washed three times in ice-cold 300 mM sucrose. The pAY9/pAY11 plasmids and pTSN2 vector were co-transformed by electroporation into the electrocompetent recipient strain and then recovered in 1 mL LB at 37 °C shaking for 1.5 hour. Successful transformants were confirmed by colony PCR using pTN7_Ver3/4 and sanger sequencing, stored at −80 °C in 25 % (v/v) glycerol.

**Construction of Transcriptional Reporter Plasmids**

The transcriptional reporter plasmids used in this study $P_{pilA}$::*yfp*, $P_{pilB}$::*yfp*, and $P_{pilMNOPQ}$::*yfp* were constructed using the established PaQa dual-reporter vector that has two promoter regions: the promoter of the gene of interest fused to yfp and the constitutive $P_{rpoD}$ promoter of the housekeeping gene *rpoD* fused to *mKate*2 as control[3]. The vector was linearized by digestion with XhoI and BsiWI. The native *pilA*, *pilB*, and *pilMNOPQ* promoter regions and *yfp* coding sequence were PCR-amplified as separate fragments using primers PpilA_F1.For/Rev (yfp) and PpilA_F2.For/Rev (pilA promoter), PpilB_F1.For/Rev (yfp) and PpilB_F2.For/Rev (pilB promoter), PpilMNOP_F1.For/Rev (*yfp*) and PpilMNOPQ_F2.For/Rev (pilMNOPQ promoter). All fragments were assembled using NEBuilder HiFi DNA Assembly Master Mix (New England Biolabs) according to the manufacturer's protocol. The products were electroporated into electrocompetent *E. coli* S17, and recovered in 1 mL LB and grown at 37 °C with shaking for 1–1.5 h. Cells were then concentrated by centrifugation, resuspended in 100 µL LB, and plated on LB agar supplemented with 100 µg/mL carbenicillin. Plates were incubated overnight at 37°C. Successful assembly was confirmed by colony PCR using primers PpilA_Ver1/2 (for *pilA* promoter), PpilB_Ver1/2 (for *pilB* promoter) and PpilMNOP_Ver1/2 (for *pilMNOPQ* promoter) which flank the insertion site and produce a diagnostic band shift. Verified clones were grown overnight in LB + 100 µg/ml carbenicillin at 37°C with shaking, mixed 1:1 with 50% (v/v) glycerol, and stored as frozen stocks at −80 °C. The respective reporter plasmid was then isolated from E. coli S17 using a Qiagen miniprep kit and introduced into recipient strains using electroporation. Electrocompetent recipient cells were prepared by growing cultures overnight in LB, diluting 1:1000 into fresh LB, harvesting at early stationary phase, washing three times in ice-cold 300 mM sucrose, and resuspending in 50 µl of the same buffer. For each electroporation, 50 µL recipient cells were mixed and with 100ng of donor plasmid, transferred to a chilled cuvette, electroporated, recovered in 1 mL LB at 37 °C shaking for 1–1.5 h. Cells were concentrated, resuspended in 100 µL LB, and plated on LB agar containing 300 µg/mL carbenicillin. Plates were incubated overnight at 37 °C. Successful plasmid maintenance was confirmed by colony PCR using primers PpilA_Ver1/2 (for pilA promoter), PpilB_Ver1/2 (for pilB promoter) and PpilMNOP_Ver1/2 (for pilMNOP promoter). Confirmed transformants were grown overnight in LB + 200 µg/ml carbenicillin and stored as 25 % glycerol stocks at −80 °C.

**Microscopy imaging and analysis**

For all assays, about 2 µL of cells were spread on an agarose pad, allowed to air-dry for 2-3 minutes and then fixed on 35mm glass bottom MatTek dish for imaging under a Nikon Ti2 Epifluorescence microscope with a ×100/1.45 NA Ph3 objective lens.

Pilus labelling and imaging: Cells were incubated with 35 ng µl$^{-1}$ Alexa Fluor 488 maleimide dye for 10 min on ice to stain the T4P. Excess dye was washed away by pelleting and resuspending the cells twice and in fresh low fluorescence EZ rich medium. T4P dynamics videos were recorded for 30 s each, with data acquisition and data analysis performed as previously described[4].

ATP biosensor imaging: The ATP biosensor and catalytic dead mutant strains were grown for 16 hours to stationary phase (SP) in 2 mL LB medium following single colonies inoculation from agar plates. At 16 hours, cells were washed by pelleting and resuspending in fresh LB medium 3 times. Cells were diluted 1:100 and grown for an additional 1 hr. After transfer to the agarose pad, cells were imaged in the 405 nm and 488 nm channels. The line scan function of ImageJ/Fiji was used to measure the mean fluorescence intensity of individual cells from the two channels separately after subtracting the background, which were then plotted relative to each other (405/488) as previously described[2].

Transcriptional reporter imaging: Overnight cultures from single colony inoculation were back-diluted 1:100 and grown for 0.5, 1, and 2 hour in 100 mL LB in a 250 mL Erlenmeyer flask at 37 °C incubator with shaking at 200 rpm. At specific time point, 2 µL of cells were spread on the agarose pad and imaged using phase-contrast and the 514 nm (YFP) and 561 nm (mKate2) channels and processed in ImageJ/Fiji. The line scan function of ImageJ/Fiji was used to measure the mean fluorescence intensity of individual cells after subtracting the background, which were then plotted relative to each other (YFP/mKate2).

Fluorescent fusion imaging: Overnight cultures from single colony inoculation were back-diluted 1:100 and grown for 0.5, 1, and 2 hours in 100 mL LB in a 250 mL Erlenmeyer flask at 37 °C incubator with shaking at 200 rpm. At specific time points, 2 µL of cells were spread on the agarose pad and imaged using phase-contrast and the 561 nm channel, and then processed in ImageJ/Fiji. The acquired images were thresholded in ImageJ/Fiji to isolate only the fusion protein signal. The raw integrated density was then measured and plotted.

**Nutrient Supplementation Assay**

Following 16 hours cell growth in LB broth from single colony inoculation, 200 µL of MOPS medium containing 20 % of glucose, lactate, or succinate was added to 1000 µL of stationary phase cells in spent medium (3.337 % final carbon source concentration), MOPS medium without carbon source was used as vehicle control. The cells were incubated at 37 C with shaking at 200 rpm for additional 20 minutes for the activation of the carbon source importers[5-8]. After 20 minutes incubation, the T4P of the cells were labelled for imaging as described above. As a control, we repeated this experiment in Δ*oprB* glucose transporter mutant, Δ*dctPQM* succinate transporter mutant, and Δ*lldD* lactate dehydrogenase mutant, all lacking the outer membrane selective porins responsible for the import of respective carbon source to the periplasm or for utilization of the carbon source.

**Phage infection assay**

Sixteen hour grown cultures of wild-type *Pseudomonas aeruginosa* PAO1 were supplemented with 20% glucose for 20 minutes as described above. After supplementation, the cells were infected with JBD68 phage at a multiplicity of infection (MOI) 0.1 and incubated for 2 minutes at 37 C, a modification to the step previously described[9]. The infected cells were pelleted and washed 5 times in fresh medium to completely remove unbound phages from the cell suspension. The phage-free infected cells suspension was back-diluted in 1:200 fresh LB and grown for 12 hours in 96 well plates using a Cerillo plate reader at 37 C, with OD readings taken at 1 hour interval. As a control, the same process was repeated for *pilA*, WT (uninfected), and WT (without glucose supplementation). Data analysis and statistical tests were done using Igor pro (64 bit).

**Biofilm dispersal assay**

The biofilm assay was done using the Nunc-immuno$^{TM}$ TSP 96 peg lid cover in 96 well microtiter plates with modification to the previously described protocol[10, 11]. Single colonies from WT PAO1 was inoculated in 2 mL of fresh LB broth and 150 µL of the inoculated LB broth was dispensed into each well of the 96 well plates. The plates were covered with Nunc-immuno$^{TM}$ TSP 96 peg lid and biofilm was allowed to form for 16 hour at 37 C with shaking at 200 rpm. After 16 hour of biofilm growth, the Nunc-immuno$^{TM}$ TSP 96 peg lid was gently washed in 150 ul of filter-sterilized spent LB medium per well. The

Nunc-immuno™ TSP 96 peg lid with the biofilm was placed onto a new 96 wells microtiter plate containing 150 μL spent LB medium supplemented with 20 % glucose in each well and the no glucose spent LB medium in 96 wells microtiter plate as a control, and then incubated for additional 2 hour at 37 °C with shaking at 200 rpm for the cells to disperse from the biofilm. The residual biofilm was stained by transferring the Nunc-immuno™ TSP 96 peg lid to a new 96 well plate containing 150 μL of crystal violet and left on the bench at room temperature for 15 minutes. Excess crystal violet was removed from the Nunc-immuno™ TSP 96 peg lid by gently rinsing the peg lid with water. The residual stained biofilm matrix was solubilized by transferring the Nunc-immuno™ TSP 96 peg lid to a 96 well plate containing 150 μL of 50 % glacial acetic acid solution. The residual biofilm was quantified by measuring the OD600 using a microplate reader. As a control, the same assay was repeated for *pilA* mutant *Pseudomonas aeruginosa* PAO1.

### Supplementary Table 1: Strains used in this study.

| Strain | Description | Reference |
|---|---|---|
| ***E. coli*** | | |
| S17 | Wild-type, used for cloning and conjugation | |
| MK 311 | pEXG2 plasmid, used for cloning | [1] |
| MK 416 | pMK89, used for cloning | [12] |
| MK 315 | pTNS2 vector, used for cloning | [13] |
| ***P. aeruginosa*** | | |
| PAO1 | Wild-type | [14] |
| MK 423 | PAO1 pilA-A86C chromosomal point mutation | [12] |
| MK 552 | PAO1 pilA-A86C, pAY1 (PpilA-YFP plasmid) | This study |
| MK 553 | PAO1 pilA-A86C, pAY2 (PpilB-YFP plasmid) | This study |
| MK 558 | PAO1 pilA-A86C, pAY7 (PpilMNOPQ-YFP plasmid) | This study |
| MK 752 | PAO1 pilA-A86C, pilB::mScarlet-i3 | This study |
| MK 575 | PAO1 pilA-A86C, pAY9 (ATP biosensor) | This study |
| MK 577 | PAO1 pilA-A86C, pAY11 (catalytic mutant ATP biosensor) | This study |
| MK 136 | PAO1 pilA-A86C, pilO::mcherry | [12] |
| MK 560 | PAO1 pilA-A86C, oprB1 deletion | This study |
| MK 809 | PAO1 pilA-A86C, lldD deletion | This study |
| MK 810 | PAO1 pilA-A86C, dctPQM deletion | This study |

### Supplementary Table 2: Primers used in this study.

| Primer | Sequence (5' to 3') | Ref. |
|---|---|---|
| dctPQM_P1 | GATACAAAGCTTGACCTCGATCTCAACAGCGG | |
| dctPQM_P2 | GACGGGTCAATCGGGACGGTTCATCGTCTTGAACATTTCGTG | |
| dctPQM_P3 | CACGAAATGTTCAAGACGATGAACCGTCCCGATTGACCCGTC | |
| dctPQM_P4 | GATACAAAGCTTCGAACAGGGTCTTCACCCAG | |
| lldD_P1 | GATACAAAGCTTCGGCGTTCCCGTTCTTCTC | |
| lldD_P2 | GGCGTCTCAGGCGCCCAGTTCAGCGGAAATGATCATTGGGCT | |
| lldD_P3 | AGCCCAATGATCATTTCCGCTGAACTGGGCGCCTGAGACGCC | |
| lldD_P4 | GATACAAAGCTTGATGCCGCCGATCTTGCA | |
| oprB1_P1 | GATACAAAGCTTGTTCGCCGCCGATGAATT | |
| oprB1_P2 | CGACGATCAGAACACCGTCTGCTTGTTCTTGTACATTTCCAGCGT | |
| oprB1_P3 | ACGCTGGAAATGTACAAGAACAAGCAGACGGTGTTCTGATCGTCG | |
| oprB1_P4 | GATACAAAGCTTCATTCGCTGCCGTTCGGA | |
| YFP_F1.FOR | GTCGTCCTTGAAAAAGATCGTACGTTCTTGG | |
| YFP_F1.REV | ATGAGCAAAGGTGAAGAACTGTTCAC | |
| PpilA_F2.FOR | ACAATGAACCCCCGCTGTTGGCGGACCAGCTTTT | |
| PpilA_F2.REV | TCTTCACCTTTGCTCATGAATCTCTCCGTTGATTATGTATAGGCC | |
| PpilB_F2.FOR | ACAATGAACCCCCGCCCCAACTTGTTGGCATCCGG | |

| | | |
|---|---|---|
| PpilB_F2.REV | TCTTCACCTTTGCTCATGGGGAAGGAATCGCAGAAGG | |
| PpilMNOP_F2.FOR | ACAATGAACCCCCGCCGGCGGCGCGCA | |
| PpilMNOP_F2.REV | TCTTCACCTTTGCTCATGACCAATTCCCTATTAGCGTTCAATACT | |
| pTN7_Ver3 | TTATCTGGTTGGCCTGCAAGG | [12] |
| pTN7_Ver4 | CGATCCATTGCTGTTGACAAAGG | [12] |
| pEXG2_Ver1 | GTTGCATGGGCATAAAGTTGCC | [12] |
| pEXG2_Ver2 | CGGGTCCTCAACGACAGG | [12] |
| pilB_FTL1_CT_F1.FOR | GGAAGCATAAATGTAAAGCAGGAGACCGCCGAGATCG | |
| pilB_FTL1_CT_F1.REV | GCCCGACCCGCCGGCATCCTTGGTCACGCGGTTG | |
| pilB_FTL1_CT_F2.FOR | GGTGGTGCCGGTGGTTAACATGGGCGTGCCGGCG | |
| pilB_FTL1_CT_F2.REV | GAGTCGACCTGCAGAGCTTTTAACGATTCCGTTTTTTCCTTGT | |
| FTL1_FP.FOR | GCCGGCGGGTCGGGC | |
| FTL1_FP.REV | ACCACCGGCACCACCCG | |

186

### Supplemental References:


1. Hmelo, L.R., et al., *Precision-engineering the Pseudomonas aeruginosa genome with two-step allelic exchange.* Nature Protocols, 2015. **10**(11): p. 1820–1841.
2. Yaginuma, H., et al., *Diversity in ATP concentrations in a single bacterial cell population revealed by quantitative single-cell imaging.* Scientific Reports, 2014. **4**(1): p. 6522.
3. Persat, A., et al., *Type IV pili mechanochemically regulate virulence factors in Pseudomonas aeruginosa.* Proc Natl Acad Sci U S A, 2015. **112**(24): p. 7563–8.
4. Koch, M.D., et al., *<i>Pseudomonas aeruginosa</i> distinguishes surfaces by stiffness using retraction of type IV pili.* Proceedings of the National Academy of Sciences, 2022. **119**(20): p. e2119434119.
5. Lin, Y.-C., et al., *The Pseudomonas aeruginosa Complement of Lactate Dehydrogenases Enables Use of <span class="smallcaps smallerCapital">d</span>- and <span class="smallcaps smallerCapital">l</span>-Lactate and Metabolic Cross-Feeding.* mBio, 2018. **9**(5): p. 10.1128/mbio.00961–18.
6. Sudarsan, S., et al., *The Functional Structure of Central Carbon Metabolism in Pseudomonas putida KT2440.* Applied and Environmental Microbiology, 2014. **80**(17): p. 5292–5303.
7. Wylie, J.L., et al., *Biophysical characterization of OprB, a glucose-inducible porin of Pseudomonas aeruginosa.* J Bioenerg Biomembr, 1993. **25**(5): p. 547–56.
8. Valentini, M., N. Storelli, and K. Lapouge, *Identification of C<sub>4</sub>-Dicarboxylate Transport Systems in Pseudomonas aeruginosaPAO1.* Journal of Bacteriology, 2011. **193**(17): p. 4307–4316.
9. Taylor, V.L., et al., *Prophages block cell surface receptors to preserve their viral progeny.* Nature, 2025. **644**(8078): p. 1049–1057.
10. Wenderska, I.B., et al., *Palmitoyl-DL-carnitine is a multitarget inhibitor of Pseudomonas aeruginosa biofilm development.* Chembiochem, 2011. **12**(18): p. 2759–66.
11. Marko, V.A., et al., *Pseudomonas aeruginosa type IV minor pilins and PilY1 regulate virulence by modulating FimS-AlgR activity.* PLoS Pathog, 2018. **14**(5): p. e1007074.
12. Koch, M.D., et al., *Competitive binding of independent extension and retraction motors explains the quantitative dynamics of type IV pili.* Proceedings of the National Academy of Sciences, 2021. **118**(8): p. e2014926118.
13. Choi, K.H., et al., *A Tn7-based broad-range bacterial cloning and expression system.* Nat Methods, 2005. **2**(6): p. 443–8.
14. Jacobs, M.A., et al., *Comprehensive transposon mutant library of Pseudomonas aeruginosa.* Proc Natl Acad Sci U S A, 2003. **100**(24): p. 14339–44.
